# Sequential maturation of supramammillary glutamatergic/GABAergic co-transmission onto adult-born dentate granule cells

**DOI:** 10.64898/2026.09.28.754986

**Authors:** Yuki Hashimotodani, Kohtarou Konno, Miwako Yamasaki, Takeshi Sakaba

## Abstract

Hypothalamic supramammillary nucleus (SuM) neurons project to the dentate gyrus and co-release glutamate and GABA, thereby regulating hippocampal information processing. However, how synapses mediating glutamate/GABA co-transmission are established during development remains unclear. Here, we investigated the functional and structural maturation of SuM inputs onto adult-born granule cells (GCs). We found that most newborn GCs received GABAergic responses without detectable glutamatergic responses, whereas a smaller population exhibited glutamate/GABA co-transmission. Newborn GCs with glutamate/GABA co-transmission showed more mature anatomical and physiological properties than GABA-only neurons, with greater dendritic complexity, longer dendritic length, lower input resistance, and stronger medial perforant path inputs. Postsynaptic depolarization failed to induce synapse unsilencing in GABA-only neurons but induced robust long-term potentiation in newborn GCs with glutamatergic transmission, indicating that plasticity at SuM inputs emerges after functional glutamatergic transmission is established. Morphological analysis showed that SuM terminals apposed to newborn GCs contained both VGluT2 and VIAAT, whereas postsynaptic receptor organization remained immature. Moreover, environmental enrichment and chemogenetic activation of SuM neurons increased the proportion of newborn GCs receiving glutamate/GABA co-transmission, suggesting that SuM activity promotes the emergence of glutamatergic transmission. Together, these results suggest that SuM inputs onto newborn GCs undergo sequential maturation from initial GABAergic transmission to glutamate/GABA co-transmission, a process likely supported by SuM activity-dependent postsynaptic maturation.

**Significance:** Synaptic transmission is generally thought to be mediated by a single neurotransmitter, yet some synapses use more than one neurotransmitter. How these dual-transmitter synapses are formed during neuronal development remains poorly understood. We studied inputs from the supramammillary nucleus to adult-born dentate granule cells, a model in which synapses mature as newborn neurons integrate into hippocampal circuits. We found that these inputs first provide GABAergic transmission and later acquire functional glutamatergic transmission as newborn granule cells mature, leading to glutamate/GABA co-transmission. This transition was promoted by activity of supramammillary neurons. These findings reveal a sequential mechanism for establishing dual-transmitter synapses and provide insight into how circuit activity shapes the functional integration of newborn neurons in the adult brain.

## Introduction

The hippocampus plays essential roles in learning and memory. The dentate gyrus (DG), a subregion of the hippocampus, is one of the few brain regions in which neurogenesis persists throughout adulthood (Gage, 2000; Gould and Gross, 2002; Ming and Song, 2011; Goncalves et al., 2016a). Adult-born DG granule cells (GCs) progressively mature and become integrated into pre-existing hippocampal circuits, where they contribute to memory processing (Deng et al., 2010; Anacker and Hen, 2017; Chavan et al., 2025). Synaptic integration of newborn GCs occurs through a multistep process over several weeks after mitosis (∼4 weeks) (van Praag et al., 2002; Esposito et al., 2005). Newborn GCs first receive tonic GABAergic signaling mediated by ambient GABA, followed by synaptic GABAergic inputs from local interneurons, the first glutamatergic inputs from mossy cells, and later perforant path (PP) inputs from the entorhinal cortex (EC) (Esposito et al., 2005; Overstreet Wadiche et al., 2005; Tozuka et al., 2005; Ge et al., 2006; Mongiat et al., 2009; Deshpande et al., 2013; Chancey et al., 2014). In addition to these well-characterized inputs, rabies virus-mediated monosynaptic retrograde tracing revealed several other sources of innervation to newborn GCs, including transient inputs from mature GCs and projections from CA3 pyramidal cells, the perirhinal cortex, the subiculum, and septal cholinergic neurons (Vivar et al., 2012; Deshpande et al., 2013). Thus, newborn GCs receive a diverse array of synaptic inputs during their development. Elucidating how individual inputs are sequentially established is essential for understanding the synaptic integration and maturation of newborn GCs.

Hypothalamic supramammillary nucleus (SuM) neurons project to the DG and innervate mature GCs and local interneurons (Pan and McNaughton, 2004; Vertes, 2015; Kesner et al., 2023). The SuM–DG pathway has recently emerged as an important hypothalamic–hippocampal circuit that contributes to various brain functions, including sleep-wake states and hippocampus-dependent cognitive functions (Renouard et al., 2015; Pedersen et al., 2017; Billwiller et al., 2020; Chen et al., 2020; Li et al., 2020; Farrell et al., 2021; Li et al., 2022a; Qin et al., 2022; Luo et al., 2025; Tian et al., 2025). One unique feature of SuM projections to the DG is that SuM neurons co-release glutamate and GABA (Pedersen et al., 2017; Hashimotodani et al., 2018; Chen et al., 2020; Li et al., 2020; Ajibola et al., 2021). Whereas the developmental mechanisms underlying the formation of excitatory and inhibitory synapses have been extensively studied in the central nervous system (Hensch, 2005; Terauchi and Umemori, 2012; Nagappan-Chettiar et al., 2023), how glutamate/GABA co-transmitting synapses are established remains largely unknown. A recent study has shown that SuM input activity promotes adult hippocampal neurogenesis and maturation of newborn GCs (Li et al., 2022b). However, the precise mechanisms underlying the formation and maturation of SuM inputs onto newborn GCs remain unclear.

In this study, we investigated the development of synaptic connections between the SuM and newborn GCs. We demonstrate that SuM inputs predominantly mediate GABAergic transmission at less mature stages, whereas glutamatergic co-transmission emerges in more mature newborn GCs. Activity-dependent synapse unsilencing is not induced in newborn GCs receiving only GABAergic SuM inputs, and morphological analyses suggest that this immature transmission may, at least in part, be associated with incomplete postsynaptic maturation. Furthermore, environmental enrichment (EE) and chemogenetic activation of SuM neurons increase the proportion of newborn GCs receiving glutamate/GABA co-transmission. Together, these findings suggest that SuM inputs onto newborn GCs progressively mature from predominantly GABAergic synapses into glutamate/GABA co-transmitting synapses, a process promoted by neural activity and experience.

## Results

### Predominance of GABAergic SuM transmission onto newborn GCs

To investigate the developmental process of SuM–newborn DG GC synapse maturation, we used glutamate decarboxylase (GAD)67-GFP (line 3) transgenic mice, in which GFP is expressed in newborn GCs approximately 1−4 weeks postmitosis in the adult hippocampus (Figure 1A) (Zhao et al., 2010). AAV-DIO-ChR2(H134R)-mCherry was injected into the SuM of GAD67-GFP mice crossed with vesicular glutamate transporter 2 (VGluT2)-Cre mice to optogenetically stimulate SuM inputs in the DG (Hashimotodani et al., 2018) (Figure 1B). Four weeks after AAV injection, we prepared acute hippocampal slices and performed whole-cell patch-clamp recordings from GFP-positive newborn GCs (76 cells) at two different membrane potentials (–57 mV for EPSCs and +9 mV for IPSCs under our recording conditions) (Hashimotodani et al., 2018) (Figure 1C). Remarkably, in the majority of newborn GCs (56.6%), light stimulation of SuM inputs evoked GABAergic synaptic responses at a holding potential of +9 mV, whereas no glutamatergic synaptic responses were detected at a holding potential of −57 mV (Figures 1D and 1E). In contrast, only 19.7% of newborn GCs exhibited both EPSCs and IPSCs (Figures 1D and 1E). We verified that the IPSCs were mediated by monosynaptic transmission rather than feed-forward inhibition as they were unaffected by NBQX and D-AP5 but were completely blocked by picrotoxin (Figure 1F). We did not detect any GFP-positive newborn GCs exhibiting EPSCs only.

**Figure 1.**
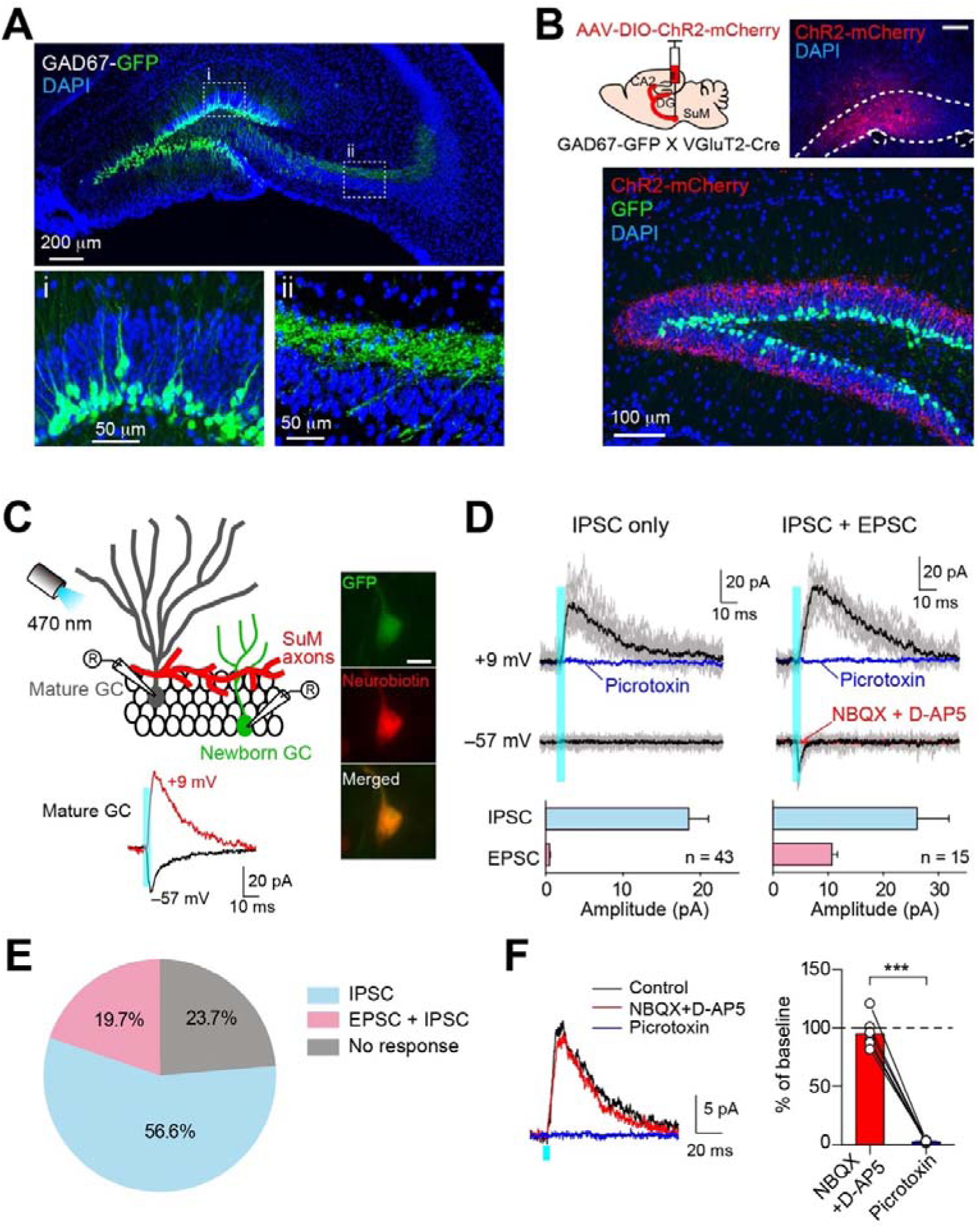
SuM inputs onto newborn GCs are predominantly GABAergic. (**A**) Fluorescence images showing GFP expression in the hippocampus of GAD67-GFP mice. Higher magnification images of the boxed region (i) and (ii) show the newborn GCs in the subgranular zone of the DG and their mossy fibers in the stratum lucidum of CA3, respectively. (**B**) (Top left) Schematic diagram showing injection of AAV-DIO-ChR2-mCherry into the SuM of GAD67-GFP mice crossed with VGluT2-Cre mice. (Top right) Fluorescence image showing the injection site of AAV in the SuM. Scale bar, 200 μm. (Bottom) ChR2(H134R)-mCherry expressing SuM axons are observed in the supragranular layer of the DG. (**C**) (Top left) Schematic of recording configuration illustrating whole-cell patch-clamp recordings from GFP-negative mature GC or GFP-positive newborn GC. ChR2(H134R)-mCherry expressing SuM axons were stimulated by 470 nm LED light illumination. (Bottom) Sample traces of EPSC (V_h_ = –57 mV, black trace) and IPSC (V_h_ = +9 mV, red trace) recorded from the mature GC. (Right) Fluorescence images of an electrophysiologically recorded, neurobiotin-filled, GFP-expressing newborn GC. Scale bar, 10 μm. (**D**) Representative traces of IPSC (top, V_h_ = +9 mV) and EPSC (middle, V_h_ = –57 mV) recorded from newborn GCs. Newborn GCs receiving IPSC only and both IPSCs and EPSCs are shown in left and right, respectively. IPSCs and EPSCs were pharmacologically verified by application of picrotoxin (100 μM) (blue) and NBQX (10 μM)/D-AP5 (50 μM) (red), respectively. (bottom) Summary graph showing the amplitude of IPSCs and EPSCs. (**E**) Pie chart showing the distribution of newborn GCs exhibiting IPSCs only, both EPSCs and IPSCs, or no response. Data are based on recordings from 76 cells. (**F**) (Left) Representative traces showing that IPSC was unaffected by NBQX/D-AP5 but was completely blocked by picrotoxin. (Right) Summary graph showing normalized IPSC amplitudes in the presence of NBQX/D-AP5 (p = 0.09, n = 8, compared with baseline, paired t-test) and picrotoxin (p < 0.001, n = 8, compared with NBQX/D-AP5, paired t-test). Data are presented as mean ± SEM. ***p < 0.001.

Similar response patterns were observed in GFP-positive newborn GCs from GAD67-GFP transgenic mice that had not been crossed with VGluT2-Cre mice (Supplementary Figure 1). In these mice, ChR2-mCherry was expressed in SuM neurons in a Cre-independent manner, excluding the possibility that VGluT2-negative SuM neurons selectively provide excitatory inputs to newborn GCs. Taken together, these results indicate that, unlike mature GCs, in which glutamate/GABA co-transmission was commonly observed (Pedersen et al., 2017; Hashimotodani et al., 2018; Billwiller et al., 2020; Chen et al., 2020; Li et al., 2020; Ajibola et al., 2021; Hirai et al., 2024), newborn GCs at approximately 1–4 weeks of age predominantly exhibited GABAergic synaptic responses without detectable glutamatergic responses.

### Newborn GCs receiving glutamate/GABA co-transmitting inputs are more mature

Interneuron-derived GABAergic inputs and cortical glutamatergic inputs onto newborn GCs in the adult DG are progressively established during neuronal maturation (Esposito et al., 2005; Overstreet Wadiche et al., 2005; Ge et al., 2006). Given that GFP-positive newborn GCs in the GAD67-GFP mice include neurons at different stages of maturation (Zhao et al., 2010), the presence of two groups of newborn GCs receiving either GABA-only transmission or glutamate/GABA co-transmission from the SuM may reflect developmental heterogeneity among the newborn GCs. To examine the relationship between SuM-evoked synaptic responses and the dendritic morphology of newborn GCs, we recorded from GFP-positive newborn GCs and subsequently performed post hoc morphological analysis. First, we filled GFP-positive and GFP-negative neurons with neurobiotin and analyzed their dendritic morphology. As expected, GFP-positive immature neurons have lower dendritic complexity and shorter total dendritic length than GFP-negative mature GCs (Figures 2A and 2B). Next, GFP-positive neurons were divided into two groups based on their SuM input-mediated synaptic responses: neurons exhibiting IPSCs only (GABA-only neurons) and neurons exhibiting both EPSCs and IPSCs (glutamate/GABA neurons). Comparison of dendritic morphology between the two groups revealed that glutamate/GABA neurons exhibited significantly greater dendritic complexity, more branches, and longer total dendritic length than GABA-only neurons (Figures 2C-2F). We also compared input resistance between glutamate/GABA neurons and GABA-only neurons as an electrophysiological indicator of maturation (Schmidt-Hieber et al., 2004; Overstreet-Wadiche et al., 2006). Glutamate/GABA neurons showed a significantly lower input resistance than GABA-only neurons (GABA-only neuron: 3.7 ± 0.5 GΩ, n = 20; glutamate/GABA neuron: 1.2 ± 0.2 GΩ, n = 12, p < 0.001, Mann–Whitney U test). Taken together, these results suggest that glutamate/GABA neurons are at a more advanced stage of maturation than GABA-only neurons.

**Figure 2.**
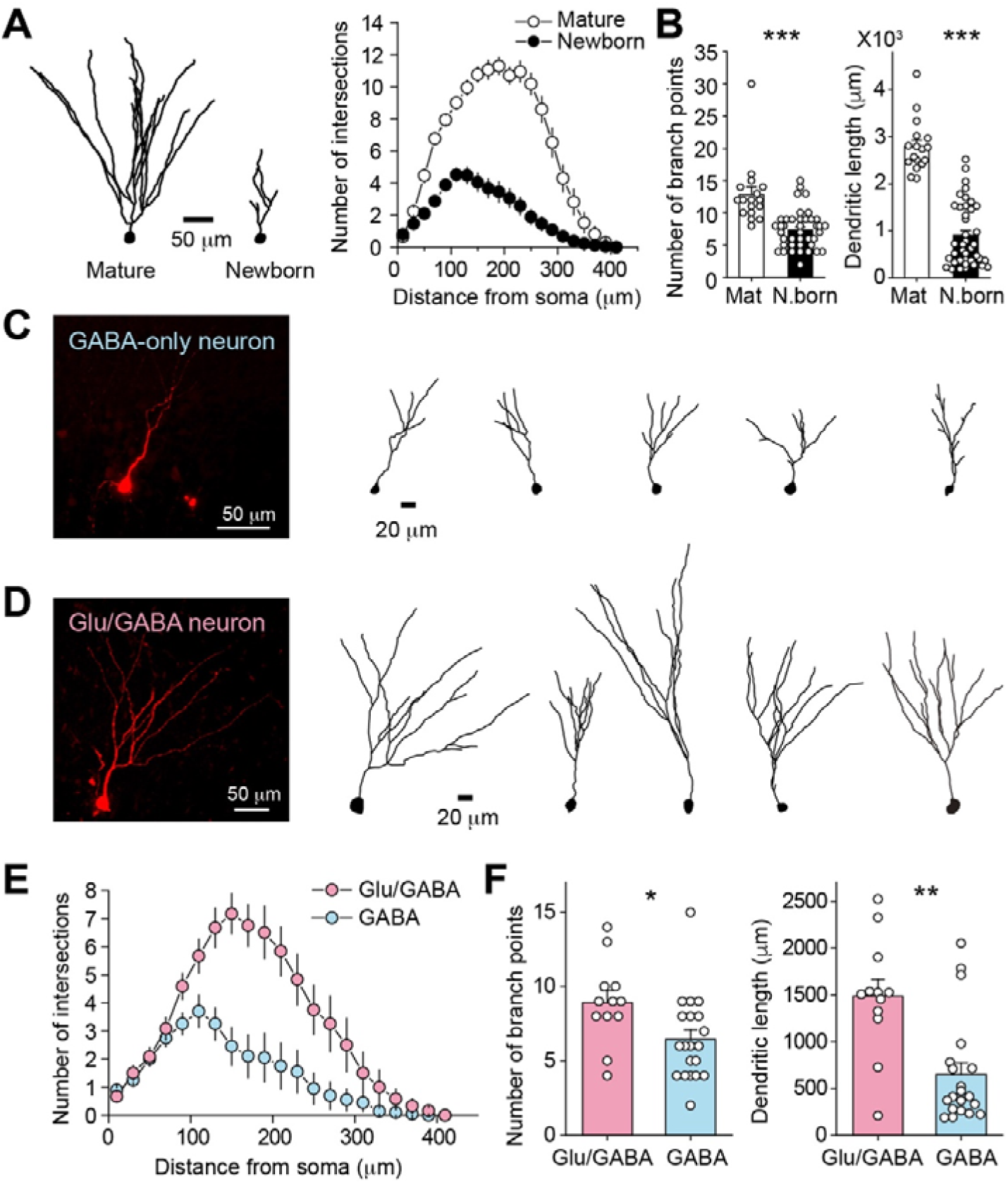
More mature newborn GCs receive glutamate/GABA co-transmitting inputs. (**A**) Example reconstructions of mature and newborn GCs (left) and Sholl analysis of dendritic complexity (right; p < 0.001, n = 17 for mature GCs and n = 40 for newborn GCs, two-way repeated measures ANOVA). (**B**) Summary graphs showing the number of branch points (left; p < 0.001, Mann–Whitney U test) and total dendritic length (right; p < 0.001, Mann-Whitney U test) in mature GCs (n = 17) and newborn GCs (n = 40). (**C, D**) Neurobiotin-filled fluorescence images of a newborn GC (left) and example reconstructions of newborn GCs (right). GABA-only neurons and glutamate/GABA neurons are shown in (C) and (D), respectively. (**E**) Sholl analysis of dendritic complexity for glutamate/GABA neurons and GABA-only neurons (p < 0.001, n = 12 for glutamate/GABA neurons and n = 20 for GABA-only neurons, two-way repeated measures ANOVA). (**F**) Summary graphs showing the number of branch points (left; p < 0.05, Mann–Whitney U test) and total dendritic length (right; p < 0.01, Mann–Whitney U test) in glutamate/GABA neurons (n = 12) and GABA-only neurons (n = 20). Data are presented as mean ± SEM. *p < 0.05, **p < 0.01, ***p < 0.001.

Previous studies have shown that newborn GCs at later developmental stages receive stronger excitatory inputs from the medial PP (MPP) (Esposito et al., 2005; Ge et al., 2006; Mongiat et al., 2009). Given that glutamate/GABA neurons are more mature than GABA-only neurons, MPP inputs may preferentially target glutamate/GABA neurons. To test this possibility, we recorded MPP-mediated EPSCs from mature GCs, glutamate/GABA neurons, and GABA-only neurons by electrically stimulating the middle molecular layer (MML) (Figures 3A and 3B). Consistent with previous studies (Mongiat et al., 2009; Kumamoto et al., 2012; Woods et al., 2018), mature GCs received more robust MPP inputs than newborn GCs (Figure 3C). Among newborn GCs, glutamate/GABA neurons received stronger MPP inputs than GABA-only neurons (Figure 3C). These results further support the idea that glutamate/GABA neurons are more mature than GABA-only neurons and suggest that the functional development of SuM inputs is associated with MPP input maturation during newborn GC maturation.

**Figure 3.**
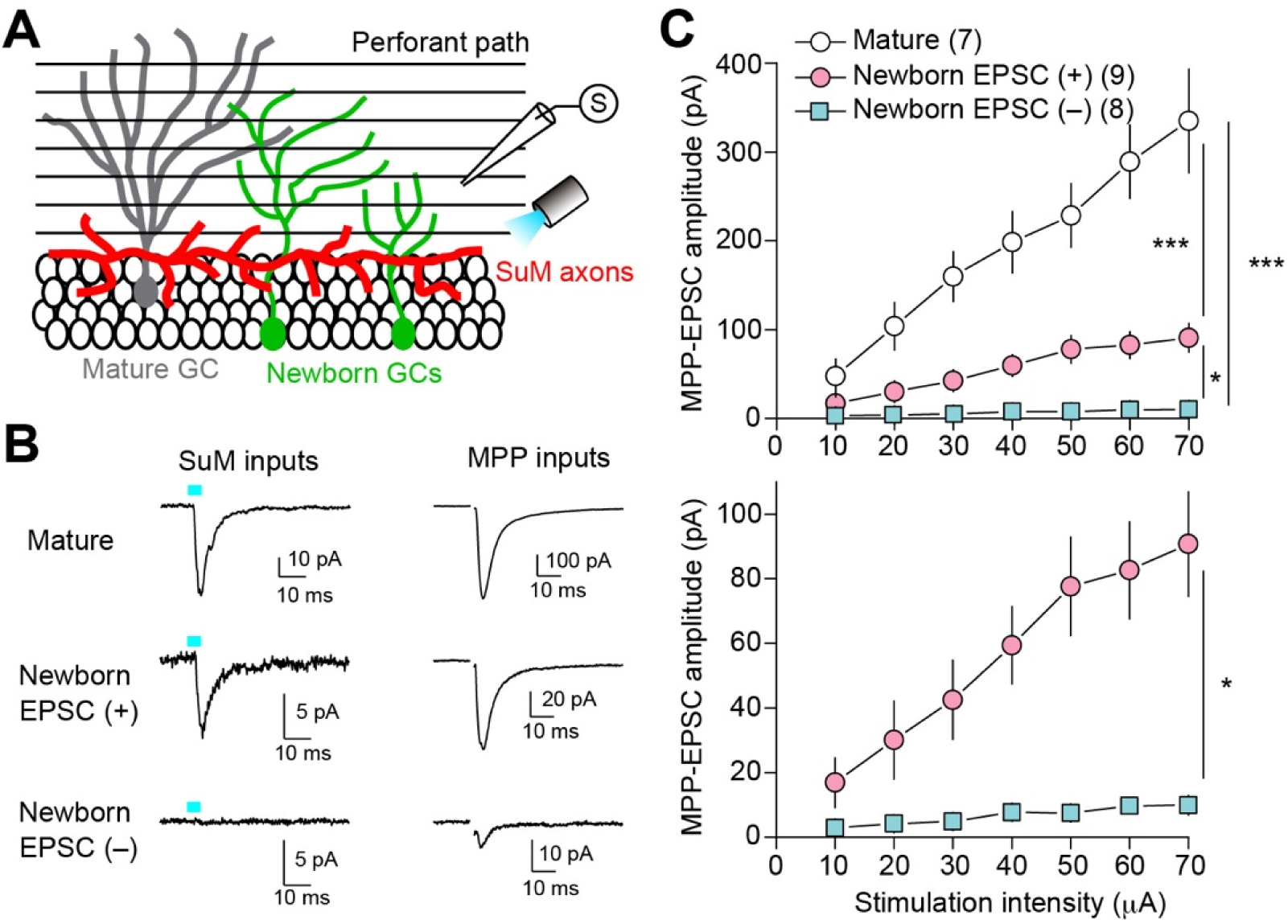
Association of SuM input maturation with MPP input maturation in newborn GCs. (**A**) Schematic diagram showing experimental configuration. A stimulating electrode was placed in the MML to activate MPP, and a blue light was delivered to activate ChR2-mCherry-expressing SuM axons. (**B**) Representative traces showing EPSCs of SuM inputs (left) and MPP inputs (right) in the mature GC (top), newborn EPSC (+) (middle, glutamate/GABA neuron), and newborn EPSC (–) (bottom, GABA-only neuron). (**C**) (top) Summary graph of MPP-EPSC amplitude plotted against the stimulation intensity. (bottom) Expanded comparison of newborn EPSC (+) and newborn EPSC (–). Two-way repeated measures ANOVA, p < 0.01, n = 7−9, Tukey’s post hoc test: mature vs newborn EPSC (+), ***p < 0.001, mature vs newborn EPSC (–), ***p < 0.001, newborn EPSC (+) vs newborn EPSC (–), *p < 0.05. Data are presented as mean ± SEM. *p < 0.05, ***p < 0.001.

### Developmental stage-dependent synaptic plasticity at SuM inputs onto newborn GCs

It has been reported that, at an early developmental stage of newborn GCs, glutamatergic transmission at PP inputs and mossy cell inputs is mediated by NMDA receptor (NMDAR)-only silent synapses (Chancey et al., 2013; Chancey et al., 2014; Li et al., 2017), and that NMDAR activation mediates synapse unsilencing (Chancey et al., 2013). To determine whether NMDAR-mediated EPSCs are present at SuM synapses onto GABA-only neurons, we recorded optogenetically evoked responses at +40 mV in the presence of GABA_A_ receptor (GABA_A_R) blocker picrotoxin. We did not detect NMDAR-mediated currents in GABA-only neurons (Figure 4A), suggesting that SuM inputs onto GABA-only neurons lack functional NMDARs. We previously reported that SuM–mature GC synapses exhibit two distinct forms of long-term potentiation (LTP): an NMDAR-dependent form (Hirai et al., 2022) and an NMDAR-independent form (Tabuchi et al., 2022). Given that SuM synapses onto GABA-only neurons lack functional NMDARs, we instead tested whether postsynaptic repetitive depolarization, which induces NMDAR-independent LTP (depol-eLTP) at SuM–mature GC synapses (Tabuchi et al., 2022), could trigger synaptic incorporation of AMPA receptors (AMPARs) and thereby lead to synapse unsilencing. Consistent with our previous report, postsynaptic depolarization induced robust LTP at SuM–GC synapses in mature GCs (Figure 4B; 193 ± 33% of baseline, n = 5, p < 0.05, paired t-test). In contrast, the same postsynaptic depolarization failed to induce synapse unsilencing in GABA-only neurons (Figure 4C; 104 ± 5% of baseline, n = 12, p = 0.18, paired t-test). In glutamate/GABA neurons, however, postsynaptic depolarization induced depol-eLTP (Figure 4D; 207 ± 37% of baseline, n = 5, p < 0.05, paired t-test). Together, these results suggest that depol-eLTP induction does not trigger the emergence of glutamatergic transmission in GABA-only neurons, and that depol-eLTP occurs at SuM–GC synapses in newborn GCs only after functional glutamatergic transmission has been established.

**Figure 4.**
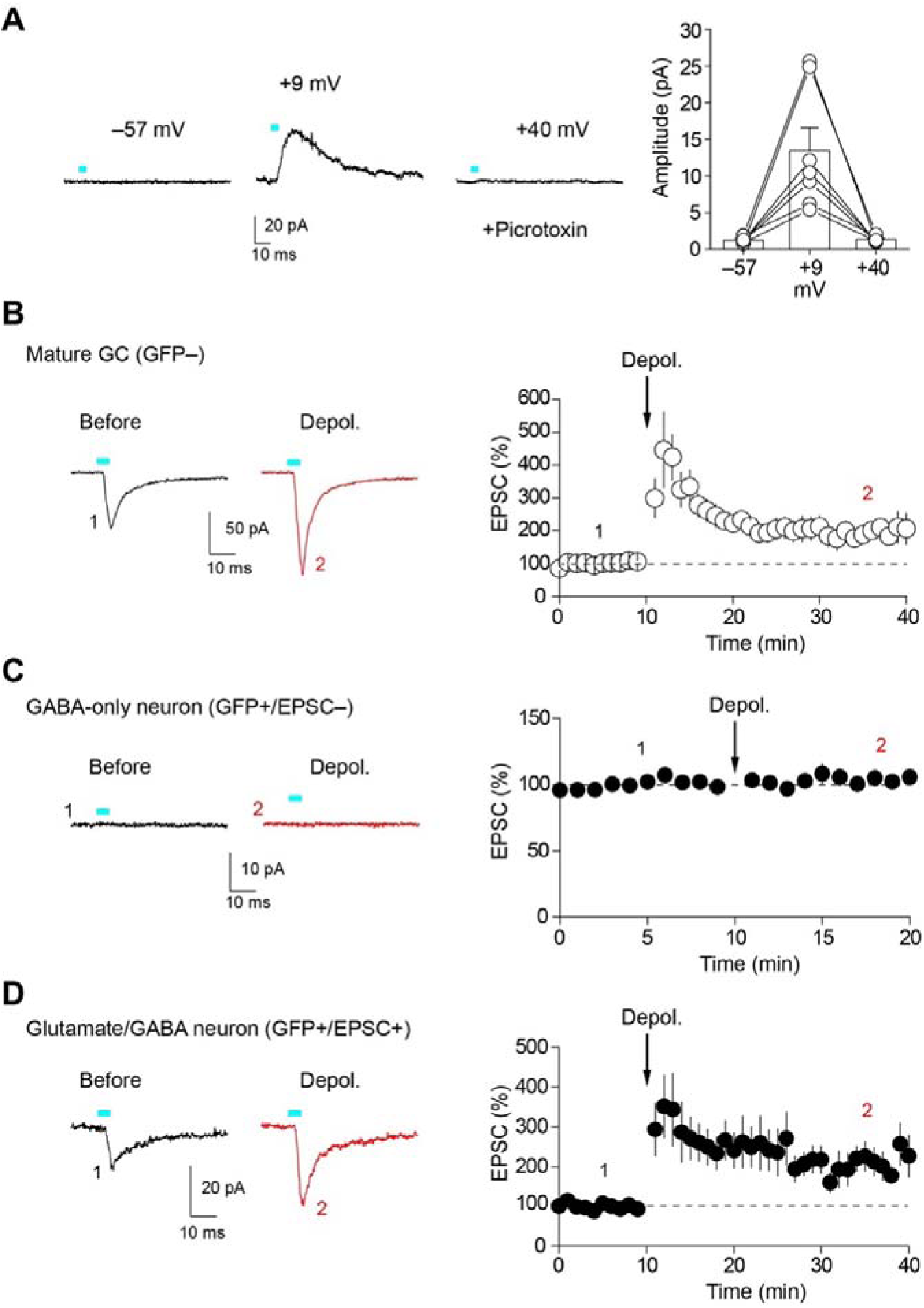
Developmental stage-dependent depol-eLTP induction at SuM inputs onto newborn GCs. (**A**) (Left) Representative traces recorded from a GABA-only neuron at holding potentials of –57 mV, +9 mV, and +40 mV. (Right) Summary plot of the amplitudes of SuM-newborn GC synaptic responses at three different holding membrane potentials (n = 7). (**B**) Representative average traces (left) and time-course summary plot (right) showing that postsynaptic depolarization induced depol-eLTP at SuM–mature GC synapses. (**C**) Representative average traces (left) and time-course summary plot (right) showing that postsynaptic depolarization failed to induce the emergence of glutamatergic synaptic responses in GABA-only neurons. Note that, because no synaptic responses were detected, the normalized baseline amplitudes were calculated using the noise level as the amplitude. (**D**) Representative average traces (left) and time-course summary plot (right) showing that postsynaptic depolarization induced depol-eLTP in glutamate/GABA neurons. Data are presented as mean ± SEM.

### SuM terminals onto newborn GCs contain both VGluT2 and VIAAT, before detectable postsynaptic receptor accumulation

The lack of detectable SuM glutamatergic transmission in early-stage newborn GCs raises the question of whether presynaptic or postsynaptic elements are immature. To examine these possibilities, we performed immunohistochemical analysis for presynaptic vesicular glutamate/GABA transporters and postsynaptic glutamate/GABA receptors. We first examined the distribution of the NMDAR subunit GluN1 at SuM-newborn GC synapses. We performed quadruple immunofluorescent labeling for GluN1 (blue), mCherry as a SuM axon marker (red), GFP as a newborn GC marker (green), and calbindin as a mature GC marker (white) (Figure 5A-5C). When we focused on SuM boutons adjacent to newborn GCs, GluN1 labeling was not detected (Figure 5C). In contrast, GluN1 was detected in mature GCs adjacent to SuM boutons (Figure 5B), as previously reported (Hirai et al., 2024). We also examined the AMPAR subunit GluA2 and found that GluA2 was not labeled in newborn GCs adjacent to SuM boutons (Figure 5D). Next, similar quadruple immunofluorescence analysis was performed for the GABA_A_R subunit GABA_A_γ2. Unexpectedly, GABA_A_γ2 immunoreactivity was hardly detectable in newborn GCs adjacent to SuM boutons, whereas it was detected in mature GCs adjacent to SuM boutons (Figure 5E-5G). These results suggest that prominent postsynaptic receptor accumulation of glutamate receptor subunit is not yet detectable at SuM–newborn GC synapses. Notably, GABA_A_γ2 accumulation was also below the detection level of our immunohistochemical analysis, despite the frequent detection of GABAergic transmission in newborn GCs.

**Figure 5.**
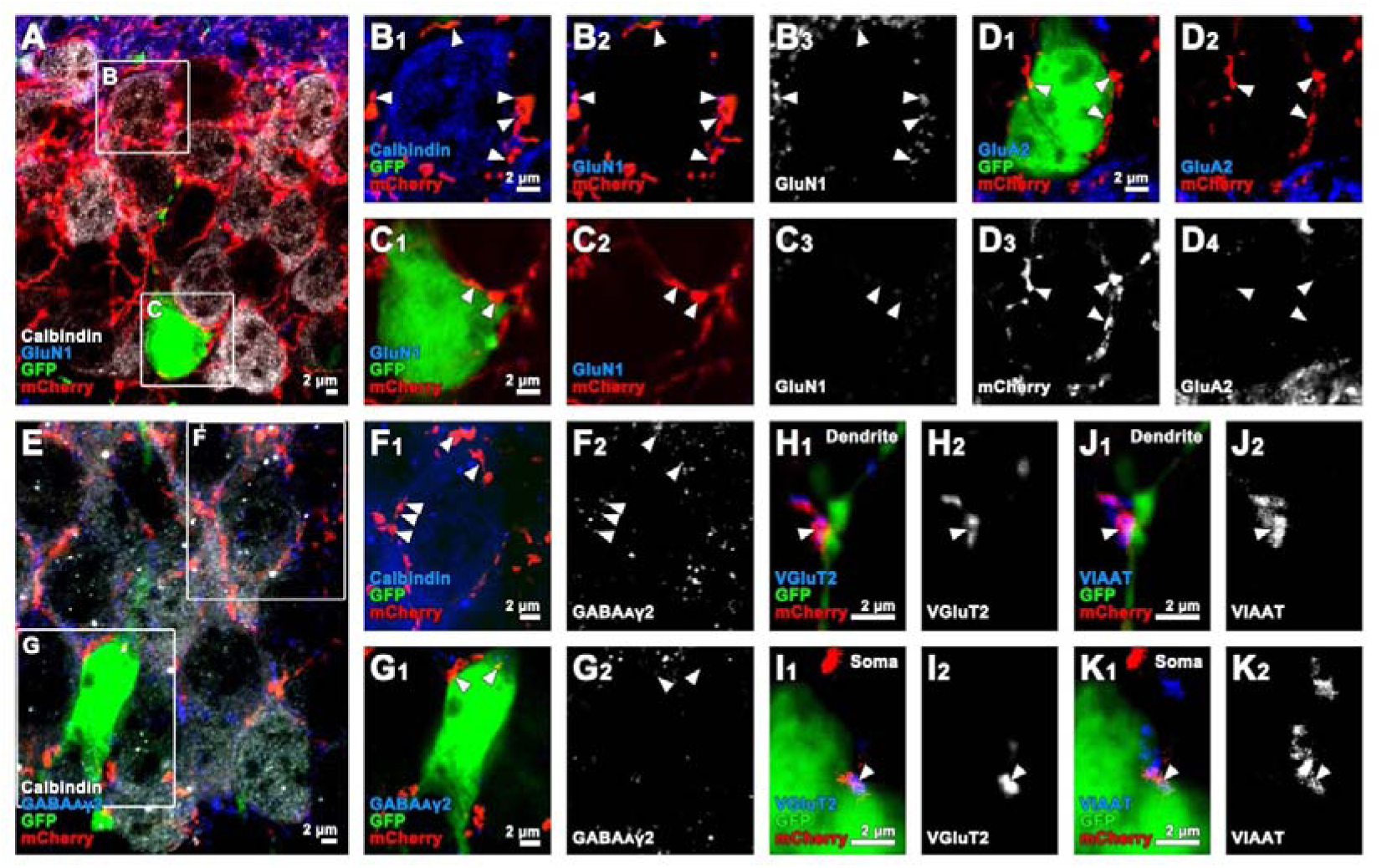
Absence of detectable postsynaptic receptors and colocalization of presynaptic VGluT2 and VIAAT at SuM-newborn GC synapses. (**A-C**) Quadruple immunofluorescent labeling for calbindin, GluN1, GFP, and mCherry. (B, C) Higher-magnification images of the boxed area in (A), showing GluN1 puncta in a calbindin-positive mature GC soma adjacent to mCherry-positive SuM terminals (B), but not in a GFP-positive newborn GC soma adjacent to SuM terminals (C). (**D**) Triple immunofluorescent labeling for GluA2, GFP, and mCherry shows no detectable GluA2 labeling in a GFP-positive newborn GC soma adjacent to mCherry-positive SuM terminals. (**E-G**) Quadruple immunofluorescent labeling for calbindin, GABA_A_γ2, GFP, and mCherry. (F, G) Higher-magnification images of the boxed area in (E), showing GABA_A_γ2 puncta in a calbindin-positive mature GC soma adjacent to mCherry-positive SuM terminals (F), but not in a GFP-positive newborn GC soma adjacent to SuM terminals (G). (**H-K**) Quadruple immunofluorescent labeling for VGluT2, VIAAT, GFP, and mCherry. VGluT2 and VIAAT are co-labeled in mCherry-positive SuM terminals adjacent to a GFP-positive newborn GC dendrite (H, J) and soma (I, K).

We next assessed the maturation of presynaptic elements by labeling VGluT2 and vesicular GABA transporter (VGAT; also known as vesicular inhibitory amino acid transporter, VIAAT) in SuM boutons onto newborn GCs (Figure 5H-5K). Colocalized labeling of VGluT2 and VIAAT was observed in mCherry-positive SuM boutons apposed to both the dendrite (Figure 5H and 5J) and soma (Figure 5I and 5K) of newborn GCs. These findings suggest that SuM terminals acquire both glutamatergic and GABAergic vesicular transporters before postsynaptic maturation.

### Maturation of glutamate/GABA co-transmitting synapses is promoted by EE and neural activity

Experience-dependent neural activity, such as exposure to EE or voluntary exercise, promotes adult neurogenesis and the synaptic integration of newborn cells (Kempermann et al., 1997; van Praag et al., 1999; Tashiro et al., 2007; Bergami et al., 2015; Alvarez et al., 2016). We examined whether exposure to EE promotes the formation of glutamate/GABA co-transmitting SuM synapses onto newborn GCs. We exposed AAV-injected adult mice to a four-week period of EE (Figure 6A) and examined the distribution of newborn GCs receiving GABA-only or glutamate/GABA co-transmitting inputs from the SuM. Consistent with previous studies (Kempermann et al., 1997; Tashiro et al., 2007; Chancey et al., 2013), exposure to EE increased the number of GFP-positive newborn GCs (Figure 6B). Remarkably, EE significantly increased the proportion of newborn GCs receiving glutamate/GABA co-transmitting inputs (Figure 6C). However, the amplitudes of both EPSCs and IPSCs were comparable between the control and EE groups (Supplementary Figure 2A), suggesting that EE promotes the emergence of glutamate/GABA co-transmitting inputs rather than enhancing the strength of individual SuM synaptic responses. Because EE increased the number of newborn GCs, it was possible that we had inadvertently recorded from more mature GFP-positive cells in the EE group, thereby contributing to the increased proportion of newborn GCs receiving glutamate/GABA co-transmitting inputs. To address this possibility, we filled a subset of recorded newborn GCs with neurobiotin and performed post hoc morphological reconstruction of these cells. However, the morphological properties of the reconstructed cells did not differ significantly between the control and EE (Supplementary Figure 2B and 2C), arguing against the possibility that the observed increase was attributable to a sampling bias toward more mature newborn GCs in the EE mice. These findings also suggest that the EE-induced promotion of glutamate/GABA co-transmission is not associated with dendritic maturation of GFP-expressing newborn GCs. Interestingly, when we focused on newborn GCs receiving glutamate/GABA co-transmitting inputs, a small but significant morphological difference was observed between the control and EE groups. The total number of branch points and total dendritic length tended to be lower in newborn GCs receiving glutamate/GABA co-transmitting inputs in the EE mice than in those in the control mice, although the differences were not statistically significant (Figure 6D).

**Figure 6.**
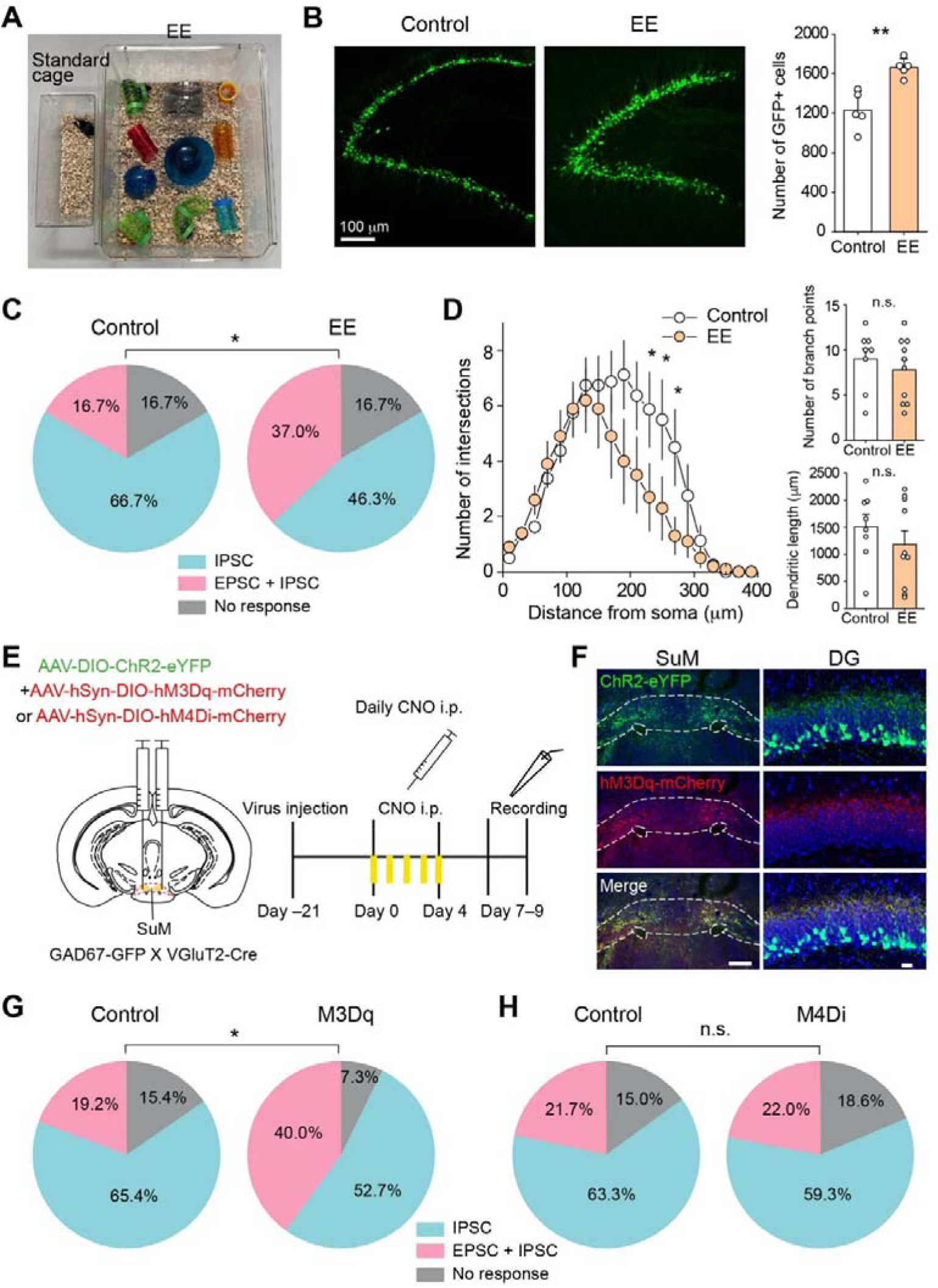
EE- and SuM neural activity-dependent promotion of the maturation of SuM inputs onto newborn GCs. (**A**) Pictures showing a standard cage (left) and an EE cage (right). (**B**) Fluorescence images of newborn GCs from control (left) and EE exposed (middle) mice. (right) Summary graph showing the EE-induced increase in the number of GFP-positive GCs (n = 5 mice, p < 0.01, unpaired t-test). (**C**) Pie charts showing the distribution of response types in newborn GCs under control (left) and EE (right) conditions (control, n = 60 cells; EE, n = 54 cells, p < 0.05, χ^2^ test). (**D**) (left) Sholl analysis of dendritic complexity of newborn GCs receiving glutamate/GABA co-transmitting inputs from the SuM in control and EE mice (control, n = 8 cells; EE, n = 10 cells, p < 0.05 at radial distances of 230, 250, and 270 μm from the soma, two-way repeated measures ANOVA followed by Tukey’s post hoc test). Summary graphs showing the number of branch points (top right; p = 0.48, unpaired t-test) and total dendritic length (bottom right; p = 0.36, unpaired t-test) in the control and EE. (**E**) Schematic diagram showing co-injection of AAV-DIO-ChR2-eYFP and AAV-hSyn-DIO-hM3Dq (or hM4Di)-mCherry into the SuM of GAD67-GFP mice crossed with VGluT2-Cre mice. Three weeks after AAV injection, CNO (1 mg/kg) was administered intraperitoneally once daily for five consecutive days. Control mice were injected with AAV-DIO-ChR2-eYFP alone and received CNO administration. (**F**) Fluorescence images showing the AAV injection site in the SuM (left) and SuM axonal projections in the DG (right). Note that both GFP-positive newborn GCs and ChR2-eYFP-expressing SuM axons exhibited green fluorescence; however, newborn GC somata could be readily identified during patch-clamp recordings. Scale bars, 200 μm (left) and 20 μm (right). (**G, H**) Pie charts showing the distribution of response types in newborn GCs under control and M3Dq (G) or under control and M4Di (H) (G: control, n = 52 cells; M3Dq, n = 55 cells, p < 0.05, χ^2^ test; H: control, n = 60 cells; M4Di, n = 59 cells, p = 0.85, χ^2^ test). Data are presented as mean ± SEM. *p < 0.05, **p < 0.01. n.s., not significant.

Sholl analysis, however, revealed significantly fewer dendritic intersections in the EE mice (Figure 6D). These results suggest that although GFP-positive newborn GCs were at similar developmental stage in the control and EE mice, exposure to EE promoted the synaptic maturation of SuM inputs, thereby allowing newborn GCs to receive glutamate/GABA co-transmitting inputs at an earlier developmental stage.

SuM neuronal activity has been reported to increase markedly in response to EE (Li et al., 2022b). We therefore hypothesized that increased SuM activity would promote the synaptic maturation of SuM inputs onto newborn GCs. To test this possibility, we used a chemogenetic approach to activate SuM neurons. We co-injected AAV-DIO-ChR2-eYFP and AAV-DIO-hM3Dq-mCherry into the SuM of GAD67-GFP mice crossed with VGluT2-Cre mice. Three weeks after AAV injection, clozapine-N-oxide (CNO) was administered intraperitoneally once daily for five consecutive days to activate SuM neurons (Figures 6E and 6F). We then prepared acute hippocampal slices from these mice and recorded optogenetically evoked EPSCs and IPSCs in newborn GCs by stimulating ChR2-expressing SuM axons. As expected, consistent with the effect of EE, chemogenetic activation of SuM neurons significantly increased the proportion of newborn GCs receiving glutamate/GABA co-transmitting inputs (Figure 6G). Conversely, we tested whether hM4Di-mediated inhibition of SuM neuronal activity would impede the synaptic maturation of SuM inputs onto newborn GCs. Using the same experimental protocol as that used for hM3Dq (Figure 6E), chemogenetic inhibition of SuM neurons had no effect on the distribution of newborn GCs receiving GABA-only inputs, glutamate/GABA co-transmitting inputs, or no detectable response compared with the control group (Figure 6H). We confirmed that CNO application suppressed the firing of hM4Di-expressing SuM neurons (Supplementary Figure 3). These results suggest that increased SuM neuronal activity enhances the synaptic maturation of SuM inputs onto newborn GCs, as reflected by an increased proportion of newborn GCs receiving glutamate/GABA co-transmitting inputs, whereas suppression of SuM neuronal activity had no effect on this process.

## Discussion

Accumulating evidence indicates that several synapses in the central nervous system are capable of co-releasing glutamate and GABA (Xu et al., 2022; Wallace and Sabatini, 2023; Ceballos et al., 2024). However, the developmental mechanisms underlying the formation of such dual-transmitter synapses remain largely unknown. In this study, we investigated the development of SuM inputs onto adult-born dentate GCs and found that these inputs initially preferentially mediate GABAergic transmission and progressively acquire glutamatergic co-transmission during neuronal maturation. Morphological analyses revealed that SuM terminals onto newborn GCs already contained VGluT2 and VIAAT. Moreover, EE and chemogenetic activation of SuM neurons promoted the emergence of glutamate/GABA co-transmission, suggesting that experience- and activity-dependent mechanisms drive the functional maturation of dual-transmitter synapses.

GABA-only neurons exhibited relatively simple dendritic branching, shorter total dendritic length, and higher input resistance compared with glutamate/GABA neurons. These neurons also showed absent or only weak MPP-evoked EPSCs. Based on previous studies describing the developmental properties of adult-born GCs, the anatomical and physiological features of GABA-only neurons are consistent with those of immature newborn GCs at 1–2 weeks after birth (Esposito et al., 2005; Overstreet Wadiche et al., 2005; Ge et al., 2006; Zhao et al., 2006; Zhao et al., 2010). Thus, we speculate that GABA-only neurons correspond to an earlier stage of newborn GC maturation (approximately 1–2 weeks after birth). In contrast, glutamate/GABA neurons showed more complex dendritic arborization, longer total dendritic length, and substantial MPP-mediated EPSCs. Because functional glutamatergic inputs from the EC are known to emerge in newborn GCs at approximately 3–4 weeks after birth and are associated with more mature morphological features (Esposito et al., 2005; Ge et al., 2006; Zhao et al., 2006; Mongiat et al., 2009; Zhao et al., 2010; Kumamoto et al., 2012; Woods et al., 2018), glutamate/GABA neurons in our study are likely to represent newborn GCs in their third to fourth week of age. Our findings are broadly consistent with a previous report showing that SuM inputs to newborn GCs acquire glutamate/GABA co-transmission at 28 days after birth (Li et al., 2022b).

During the development of adult-born GCs, immature GCs exhibit distinct forms of synaptic plasticity at different developmental stages (Jahn and Bergami, 2018). At an early developmental stage (approximately 1–2 weeks after birth), newborn GCs undergo synapse unsilencing, in which AMPARs are incorporated into silent NMDAR-only synapses in a manner requiring depolarizing GABA signaling and NMDAR activation, thereby establishing initial glutamatergic inputs, presumably from mossy cells (Chancey et al., 2013; Chancey et al., 2014). At a subsequent developmental stage, immature GCs exhibit high intrinsic excitability and a lower threshold for LTP induction of PP inputs at approximately 4–6 weeks after birth (Schmidt-Hieber et al., 2004; Ge et al., 2007b; Kennedy et al., 2024). We found that glutamate/GABA neurons, whose developmental stage may partially overlap with the period of enhanced plasticity, exhibited depol-eLTP at SuM synapses. In contrast, GABA-only neurons appeared to represent an early developmental stage comparable to that associated with synapse unsilencing at mossy cell inputs, yet postsynaptic depolarization failed to induce synaptic plasticity at SuM inputs onto these cells. We previously demonstrated that the same postsynaptic depolarization induces synapse unsilencing at SuM inputs onto mature GCs (Tabuchi et al., 2022). Therefore, the lack of synapse unsilencing at SuM inputs onto GABA-only neurons may be partly explained by immature postsynaptic specialization for glutamatergic transmission at these synapses. This interpretation is supported by our morphological analysis showing that postsynaptic AMPARs and NMDARs were not detectable at SuM-newborn GC synapses. Alternatively, SuM synapses may undergo synapse unsilencing during a brief critical period that was not captured by our recordings. Future studies using temporally precise approaches, such as retroviral labeling of newborn GCs (Enikolopov et al., 2015), will be necessary to determine when and how functional glutamatergic transmission emerges at SuM inputs during newborn GC maturation.

Previous anatomical studies have shown that the vast majority of SuM boutons in the DG are co-labeled with VGluT2 and VIAAT (Boulland et al., 2009; Soussi et al., 2010; Root et al., 2018; Billwiller et al., 2020; Ajibola et al., 2021; Hirai et al., 2024). However, whether this dual vesicular-transporter phenotype is already present in SuM terminals contacting newborn GCs has remained unknown. In the present study, we demonstrate for the first time that VGluT2 and VIAAT are colocalized within the same SuM terminals apposed to immature GCs. If SuM terminals have already acquired both VGluT2 and VIAAT despite the lack of EPSCs in GABA-only neurons, an important question is how these SuM-derived presynaptic terminals are established onto newborn GCs. Previous electron microscopy analysis has shown that dendritic filopodia of newborn GCs preferentially contact axonal boutons that already form synapses onto mature GCs, suggesting that immature GCs initially establish synaptic contacts with preexisting presynaptic boutons, rather than through de novo bouton formation (Toni et al., 2007). Based on this model, SuM-derived boutons that already form synapses with mature GCs may serve as presynaptic partners for newborn GCs and contribute to the initial formation of SuM inputs onto these cells. Together with our finding that SuM terminals onto early-stage newborn GCs already contain both VGluT2 and VIAAT, this model suggests that the emergence of glutamate/GABA co-transmission may depend largely on the maturation of postsynaptic organization rather than on the acquisition of presynaptic transmitter machinery.

Experience is known to strongly influence the maturation of adult-born GCs. In particular, EE promotes dendritic elongation (Alvarez et al., 2016; Goncalves et al., 2016b) and contributes to the reorganization of presynaptic connectivity in newborn GCs (Bergami et al., 2015). In the present study, we found that EE increased the proportion of newborn GCs receiving glutamatergic transmission from SuM inputs. Notably, however, glutamate/GABA neurons in the EE group did not show enhanced dendritic maturation; rather, their dendrites were less complex than those in the control group. These findings suggest that EE promotes the emergence of glutamatergic co-transmission at SuM inputs independently of gross dendritic maturation of newborn GCs. It should be noted that, in the GAD67-GFP mice used in this study, GFP is transiently expressed in newborn GCs during a limited developmental window, approximately 1–4 weeks after birth. Therefore, we cannot exclude the possibility that a subset of newborn GCs underwent accelerated maturation in response to EE and consequently downregulated GFP expression, thereby being excluded from our recordings and morphological analyses.

At the early phase of synaptic integration of newborn GCs, excitatory action of tonic and phasic GABA signaling due to high intracellular Cl^-^ concentration contributes to the dendritic maturation and synaptic integration of immature GCs (Overstreet Wadiche et al., 2005; Tozuka et al., 2005; Ge et al., 2006; Overstreet-Wadiche et al., 2006; Markwardt et al., 2011; Chancey et al., 2013; Heigele et al., 2016). Given that SuM neurons show increased activity following EE exposure (Li et al., 2022b), EE-induced promotion of glutamate/GABA co-transmission may be attributed to the depolarizing action of GABAergic signaling from the SuM. Consistent with this idea, chemogenetic activation of SuM neurons, which likely increased SuM-derived GABA release, also enhanced the emergence of glutamate/GABA co-transmitting inputs. GABAergic depolarization is expected to trigger several Ca^2+^-dependent intracellular cascades involved in adult neurogenesis and the development of newborn GCs (Ge et al., 2007a). Cyclic AMP response element-binding protein (CREB) signaling has been implicated as a downstream mechanism linking GABAergic depolarization to the survival and development of newborn GCs (Jagasia et al., 2009). Interestingly, CREB phosphorylation has been observed in newborn GCs during the period of depolarizing GABA action (Overstreet-Wadiche et al., 2006; Jagasia et al., 2009), raising the possibility that CREB signaling contributes to the functional maturation of glutamatergic synapses formed by SuM inputs onto newborn GCs. In contrast to SuM activation, chemogenetic inhibition of SuM neurons did not alter the distribution of response types in newborn GCs. One possible explanation is that other sources of tonic and phasic GABAergic signaling compensated for the reduction in SuM-derived GABAergic input. These findings suggest that the basal establishment of SuM inputs onto newborn GCs may proceed, at least in part, independently of SuM neuronal activity, whereas increased SuM activity further facilitates this process and promotes the emergence of glutamate/GABA co-transmitting inputs.

Why are GABAergic and glutamatergic transmission established sequentially, rather than simultaneously, at SuM inputs onto newborn GCs? The transition from GABAergic transmission to glutamate/GABA co-transmission at SuM inputs may reflect the order in which newborn GCs become integrated into hippocampal circuits. In this regard, SuM inputs onto newborn GCs appear to follow a developmental sequence that parallels the established order of synaptic integration, in which early GABAergic signaling is followed by glutamatergic inputs from mossy cells and the PP (Esposito et al., 2005; Overstreet Wadiche et al., 2005; Tozuka et al., 2005; Ge et al., 2006; Mongiat et al., 2009; Deshpande et al., 2013; Chancey et al., 2014). Our results suggest that GABAergic SuM inputs promote the maturation of SuM-newborn GC synapses. Early GABAergic SuM inputs also promote the integration and maturation of immature GCs (Li et al., 2022b). These findings suggest that early GABA-only SuM inputs may play an important role in the initial phase of SuM–newborn GC circuit maturation. As development proceeds, glutamatergic co-transmission at SuM inputs emerges during the period when newborn GCs exhibit transiently enhanced excitability (Schmidt-Hieber et al., 2004; Ge et al., 2007b) and a high excitation/inhibition balance (around 4 weeks after birth), allowing them to be recruited by relatively weak afferent activity (Marin-Burgin et al., 2012). Thus, SuM glutamate/GABA co-transmission may contribute to the functional recruitment of newborn GCs during this later, functionally sensitive period. Previous studies have mainly focused on local interneurons, mossy cells, and PP inputs as key regulators of adult-born GC development (Esposito et al., 2005; Overstreet Wadiche et al., 2005; Tozuka et al., 2005; Ge et al., 2006; Mongiat et al., 2009; Deshpande et al., 2013; Chancey et al., 2014). Our findings extend this view by suggesting that long-range hypothalamic projections from the SuM also participate in the activity-dependent maturation of newborn GCs. Given that SuM-modified adult-born GCs have been implicated in memory and emotion processing (Li et al., 2022b; Li et al., 2023), elucidating how SuM inputs are formed and mature onto newborn GCs may provide important insight into the circuit mechanisms by which adult-born GCs contribute to hippocampal functions.

## Materials and methods

### Animals

VGluT2-Cre mice (Jackson Laboratory, Slc17a6^tm2(cre)Lowl^/J, stock #016963) (Vong et al., 2011) were crossed with GAD67-GFP (line 3) mice (Jackson Laboratory, Tg(Gad1-EGFP)3Gfng/J, stock #007673) (Zhao et al., 2010). Double-transgenic offspring of either sex aged 7–9 weeks were used for all experiments except for Supplementary Figure 1. All animals were group housed in a temperature- and humidity-controlled room under a 12 hr light/12 hr dark cycle. Water and food were provided *ad libitum*. The experiments were approved by the animal care and use committee of Doshisha University and Hokkaido University and were performed in accordance with the guidelines of the committees.

### Stereotaxic injections

Mice on postnatal days 19 to 21 were placed in a stereotaxic frame, and anesthetized with isoflurane (1.5−2.5%). A beveled glass capillary pipette connected to a microsyringe pump (UMP3, WPI) was used for the viral injection. 200 nL of AAV-EF1a-DIO-hChR2(H134R)-mCherry (Addgene) or AAV-EF1a-DIO-hChR2(H134R)-eYFP (Addgene) was injected into the SuM (relative to bregma, AP: −2.2 mm, ML: ±0.5 mm, DV: −4.85 mm) at a rate of 50 nL/min. The glass capillary was remained at the target site for 5 min before the beginning of the injection and was removed 10 min after infusion.

### Hippocampal slice preparation

Acute transverse hippocampal slices were prepared as described previously (Hashimotodani et al., 2017). Except for the EE and chemogenetic experiments described below, mice were decapitated under isoflurane anesthesia 4 weeks after AAV injection. Briefly, the hippocampi were isolated, embedded in an agar block, and cut at 300-μm-thick slices using a vibratome (VT1200S, Leica Microsystems) in an ice-cold cutting solution containing (in mM): 215 sucrose, 20 D-glucose, 2.5 KCl, 26 NaHCO_3_, 1.6 NaH_2_PO_4_, 1 CaCl_2_, 4 MgCl_2_, and 4 MgSO_4_. Brain blocks, including the interbrain and midbrain, were also isolated and fixed in 4% paraformaldehyde (PFA) for post hoc our internal check of the injection site (Tabuchi et al., 2022). Hippocampal slices were transferred to an incubation chamber and incubated at 33.5°C in the cutting solution. After 30 min of incubation, the cutting solution was replaced with an extracellular artificial cerebrospinal fluid (ACSF) containing (in mM): 124 NaCl, 2.5 KCl, 26 NaHCO_3_, 1 NaH_2_PO_4_, 2.5 CaCl_2_, 1.3 MgSO_4_ and 10 D-glucose at 33.5°C. The slices were stored at room temperature for at least 1 h before recording. Both the cutting solution and the ACSF were oxygenated with 95% O_2_ and 5% CO_2_. After recovery, slices were transferred to a submersion-type recording chamber for electrophysiological analysis.

### Electrophysiology

Whole-cell recordings were made from mature or newborn GCs using an EPC10 amplifier (HEKA Elektronik) or an IPA amplifier (Sutter Instruments) under infrared differential interference contrast (IR-DIC) visualization with an Olympus BX51WI microscope. The data were filtered at 2.9 kHz and sampled at 20 kHz. For voltage-clamp recordings, we used patch pipettes (3−6 MΩ) filled with an intracellular solution with the following composition (in mM): 125 Cs-gluconate, 10 CsCl, 10 HEPES, 0.2 EGTA, 2 Mg-ATP, 0.3 Na_3_GTP, 10 phosphocreatine, pH 7.3 adjusted with CsOH (295 mOsm). Consistent with previous study (Zhao et al., 2010), we confirmed that GFP-positive newborn GCs had higher input resistance than GFP-negative mature GCs (newborn: 2.59 ± 0.27 GΩ, n = 76; mature: 208.4 ± 12.0 MΩ, n = 30, p < 0.001, Mann–Whitney U test). For post hoc morphological visualization of GCs, neurobiotin (0.2%, Vector Laboratories) was included in the intracellular solution. ChR2-expresing SuM fibers were activated at 0.05 Hz by a pulse of 470 nm blue light (1–5 ms duration, 5.0–10.5 mW/mm^2^) delivered through a ×40 objective attached to a microscope using an LED (Mightex or ThorLabs). The illumination field was centered over the recorded cell. For each mouse, we first confirmed that light activation of ChR2-expressing SuM inputs evoked synaptic responses in mature GCs before performing recordings from GFP-positive newborn GCs. For extracellular stimulation of MPP inputs, a patch pipette with a broken tip (with diameter of ∼20−30 μm) filled with ACSF was placed in the MML to activate MPP inputs (100 μs pulse width, 10–70 μA). Depol-eLTP was induced by repeated postsynaptic depolarizations (2 s duration repeated 10 times every 5 s from a holding potential of −70 mV to 0 mV) as reported previously (Tabuchi et al., 2022). Recordings shown in Supplementary Figure 3 were obtained from GFP-positive neurons in the SuM under current-clamp conditions using an intracellular solution with the following composition (in mM): 136 K-gluconate, 4 KCl, 10 HEPES, 0.2 EGTA, 5 NaCl, 2 MgATP, 0.3 Na3GTP, 10 phosphocreatine, pH 7.3 adjusted with KOH (288–294 mOsm). SuM-containing coronal slices (170 μm thick) were prepared using the same protocol as that used for hippocampal slices. Input resistance was obtained from the current response to a 5-mV hyperpolarizing voltage step in voltage-clamp mode. Series resistance (8−20 MΩ) was uncompensated and monitored throughout experiments with a −5 mV, 50 ms voltage step, and cells that exhibited a significant change in the series resistance more than 20% were excluded from analysis. All experiments were performed at 30−33°C in the recording chamber perfused (2 mL/min) with oxygenated ACSF.

### Neurobiotin staining and morphological reconstructions

After the recordings, the slices were fixed overnight in 4% PFA at 4°C. The fixed slices were subsequently treated with PBS containing 0.5% Triton X-100 for 1 hours at room temperature. Slices were then incubated with streptavidin-conjugated Alexa Fluor 568 (1:500, Thermo FisherScientific) in PBS and 0.1% Triton X-100 overnight at room temperature. After washing 3 times with PBS, the slices were mounted on glass slides with VECTASHIELD antifade mounting medium (Vector Laboratories).

Fluorescence images were captured by using a fluorescence microscope (BZ-X800, Keyence), and labeled neurons were reconstructed using Neurolucida (MBF Bioscience). Sholl analysis was performed using Neurolucida by counting the number of intersecting dendrites with concentric circles, which were drawn at 20 μm increments from the center of soma. Dendritic length and branch points were also evaluated.

### Fluorescence histology

For visualizing the fluorescence images of ChR2(H134R)-eYFP, ChR2(H134R)-mCherry, and GFP-expressing newborn cells, mice under deep pentobarbital anesthesia (100 mg/kg of body weight, intraperitoneally) were perfused with 4% PFA. Brains were removed and stored in 4% PFA for 4 h at room temperature, then transferred to PBS and left overnight at 4°C. Coronal brain slices containing the SuM or the hippocampus were sectioned at 100 µm using a vibratome (VT1200S, Leica Microsystems). Sections were rinsed twice in PBS and mounted on glass slides with VECTASHIELD antifade mounting medium (Vector Laboratories). Fluorescence images were acquired using a fluorescence microscope (BZ-X800, Keyence). To quantify newborn cells, GFP-positive neurons in the DG were automatically counted using the built-in Hybrid Cell Count system of the BZ-X800. Cell counts were obtained from five animals in each of the control and EE groups. Six sections per animal, spanning the dorsal to ventral hippocampus, were used for measurement.

### Antibodies

Primary antibodies raised against the following molecules were used: calbindin (Miura et al., 2006), DsRed, GABA_A_ receptor γ2 subunit (Rovo et al., 2014), GFP (Takasaki et al., 2010), GluA2 (Yamasaki et al., 2011), GluN1 (Siegel et al., 1994), VGluT2 (Miyazaki et al., 2003), and VIAAT (Miura et al., 2006). Information on the target molecule, antigen sequence, host species, specificity, reference, NCBI GenBank or UniProt accession number, and RRID for each primary antibody is summarized in Table S1.

### Immunofluorescence

Under deep pentobarbital anesthesia (100 mg/kg, i.p.), mice were euthanized by cervical dislocation, and the brains were removed and immersed in a glyoxal fixative solution (9% glyoxal [Sigma], 8% acetic acid, pH 4.0) (Konno et al., 2023). Brains were post-fixed at 4°C in the same fixative overnight and immersed in 30% sucrose for cryoprotection. Coronal sections (50 μm thick) were prepared using a cryostat (CM1860, Leica Microsystems). Free-floating immunostaining was performed in glass test tubes. Phosphate-buffered saline containing 0.5% Triton X-100 (PBS-T, pH 7.4) was used for all incubation and washing steps. Sections were blocked with 10% normal donkey serum (Jackson ImmunoResearch, West Grove, PA, USA) for 20 min and then incubated overnight at room temperature with a mixture of primary antibodies (1 μg/ml each). After washing, sections were incubated with Alexa Fluor Plus 405-, Alexa Fluor 488-, Alexa Fluor 647-, or Cy3-conjugated species-specific anti-IgG antibodies (1:400 each; Thermo Fisher Scientific) for 2 h at room temperature. After washing, sections were air-dried and mounted with VectaShield (Vector Laboratories, Newark, CA, USA). Photographs were taken with a confocal laser microscope (FV1200, Evident, Tokyo, Japan) using a 60× objective.

### EE exposure

One week after AAV injection, mice were exposed to an EE consisting of a large cage (45 cm × 58 cm × 20 cm) equipped with colored plastic tunnels, two running wheels, and domes. Control mice were housed in a standard cage (15 cm × 35 cm × 15 cm). After four weeks of EE exposure, animals were used for electrophysiological experiments.

### Chemogenetics

For chemogenetic activation of SuM neurons, a 1:1 mixture of AAV-EF1a-DIO-hChR2(H134R)-eYFP and AAV-hSyn-DIO-hM3Dq-mCherry was bilaterally injected into the SuM (relative to bregma, AP: −2.2 mm, ML: ±0.5 mm, DV: −4.85 mm). For chemogenetic inhibition, AAV-hSyn-DIO-hM4Di-mCherry was used instead of AAV-hSyn-DIO-hM3Dq-mCherry. For the control group, AAV-EF1a-DIO-hChR2(H134R)-eYFP alone was injected into the SuM. Three weeks after AAV injection, mice were injected i.p. with CNO (1 mg/kg) once daily for five consecutive days. Control mice, which did not express hM3Dq or hM4Di, were also administered CNO to control for its potential off-target effects (Gomez et al., 2017). Three days after the final CNO administration, mice were subjected to electrophysiological experiments.

### Pharmacology

Each reagent was dissolved in water, or DMSO, depending on the manufacture’s recommendation to prepare a stock solution, and stored at −20°C. NBQX, D-AP5, and CNO were purchased from Tocris Bioscience. Picrotoxin was purchased from the Tokyo Chemical Industry. Reagents were bath applied following dilution into ACSF from the stock solutions immediately before use.

### Statistics

Statistical analyses were performed using OriginPro software (OriginLab, USA). Normality of the distributions was assessed using the Shapiro−Wilk test. For samples with normal distributions, Student’s unpaired and paired two-tailed t-tests were used to assess the between-group and within-group differences, respectively. For samples that were not normally distributed, a non-parametric paired sample Wilcoxon signed-rank test and Mann–Whitney U test were used. Differences among two or multiple samples were assessed by using one-way or two-way ANOVA followed by Tukey’s post hoc test for multiple comparisons. A *χ* ^2^ (chi-squared) test was used to compare the proportion of responsive cells following optogenetic stimulation of SuM fibers between two groups. Statistical significance was set at p < 0.05 (***, **, and * indicates p < 0.001, p < 0.01 and p < 0.05, respectively). All values are reported as the mean ± SEM.

## Acknowledgement

This work was supported by the Grants-in-Aid for Scientific Research (25H02522, and 25K02366 to Y.H.; 22H04926 to M.Y.; 24K21279 and 24K02125 to T.S.) from Japan

Society for the Promotion of Science (JSPS), the Takeda Science Foundation (to Y.H.).

## Author contributions

Y.H., K.K., M.Y., and T.S. designed the experiments. Y.H. K.K., and M.Y. performed the experiments and data analysis. Y.H. wrote the manuscript with input from all authors.

## Supplementary figures

**Supplementary figure 1.**
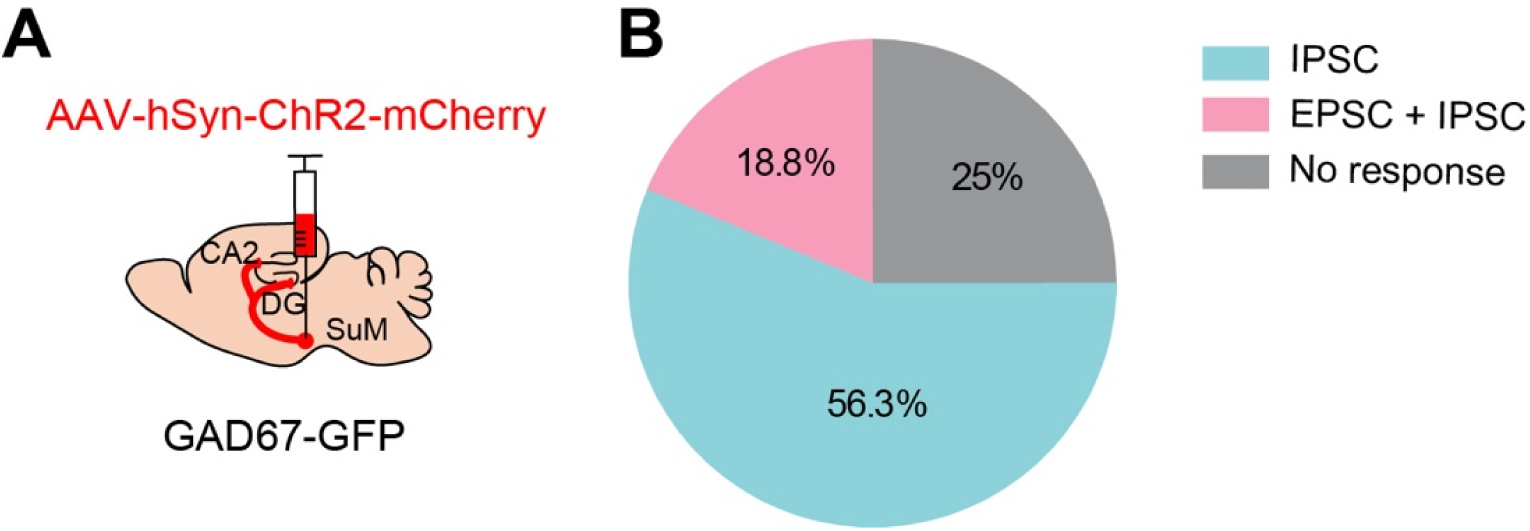
Functional characterization of SuM transmission onto newborn GCs in GAD67-GFP mice. (**A**) Schematic diagram showing injection of AAV-hSyn-ChR2-mCherry into the SuM of GAD67-GFP mice. (**B**) Pie chart showing the distribution of response types in newborn GCs. Data are based on recordings from 32 cells.

**Supplementary figure 2.**
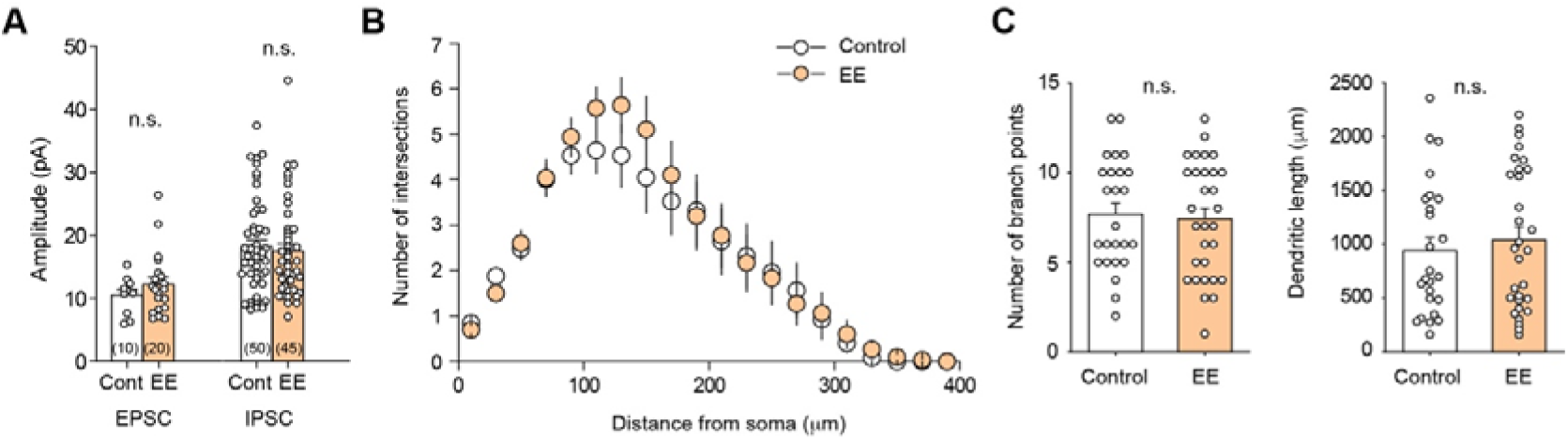
Amplitudes of EPSCs/IPSCs and dendritic morphology of newborn GCs in control and exposure to EE. (**A**) Summary graph showing the amplitudes of EPSCs and IPSCs obtained from newborn GCs in control and EE mice (EPSC: p = 0.42, Mann–Whitney U test; IPSC: p = 0.59, unpaired t-test). (**B**) Sholl analysis of dendritic complexity for control and EE mice (control, n = 25; EE, n = 30, p = 0.96, two-way repeated measures ANOVA). (**C**) Summary graphs showing the number of branch points (left; p = 0.77, unpaired t-test) and total dendritic length (right; p = 0.65, Mann–Whitney U test) in the control and EE.

**Supplementary figure 3.**
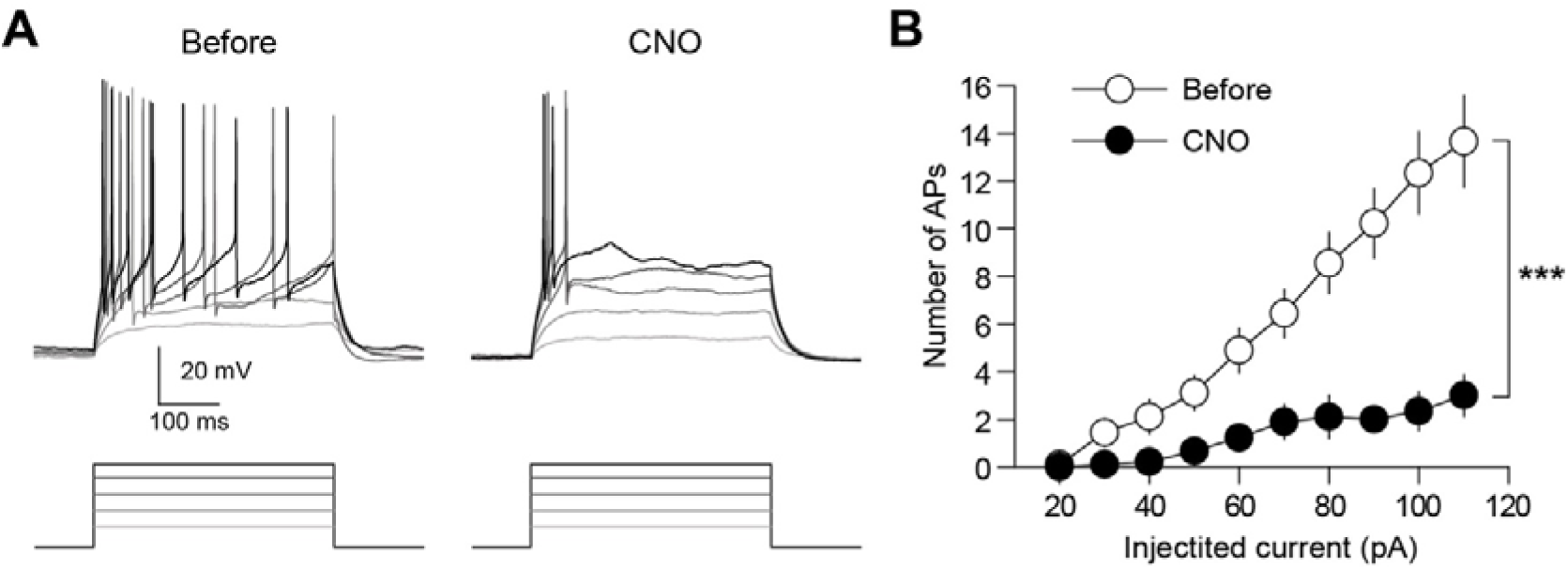
CNO suppressed hM4Di-expressing SuM neurons. (**A**) Representative traces showing the action potentials in response to current injections (20, 40, 60, 80, and 100 pA) in hM4Di-expressing SuM neurons before (left) and after application of 10 μM CNO (right). (**B**) Summary graph showing the effect of CNO on the firing of hM4Di-expressing SuM neurons (p < 0.001, n = 9, two-way repeated measures ANOVA).

**Table S1.**
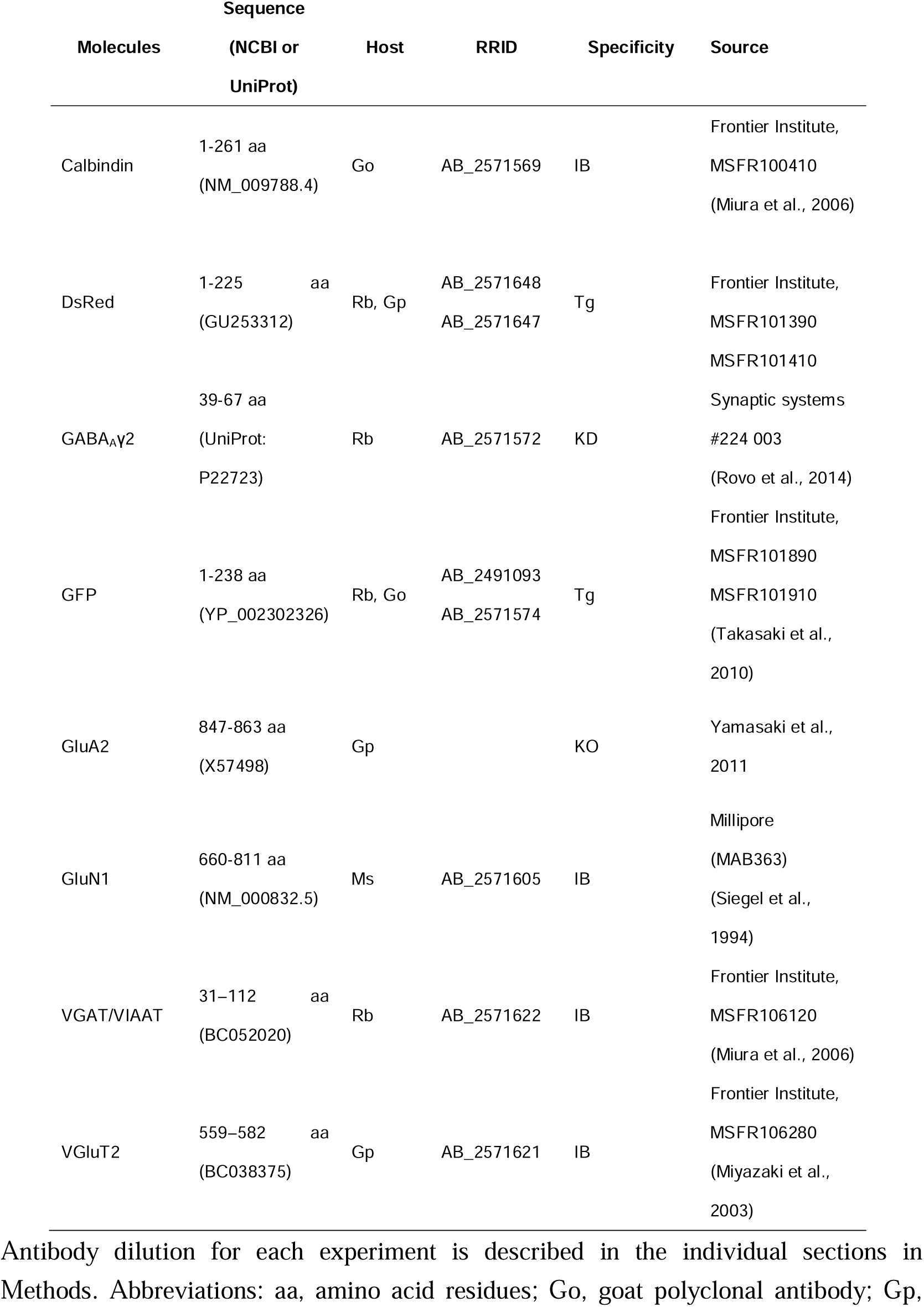
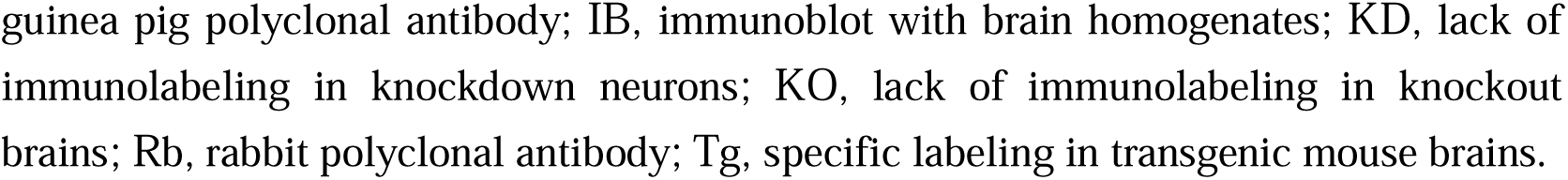
List of primary antibodies used in the present study.

| Molecules | Sequence<br>(NCBI or<br>UniProt) | Host | RRID | Specificity | Source |
| --- | --- | --- | --- | --- | --- |
| Calbindin | 1-261 aa<br>(NM_009788.4) | Go | AB_2571569 | IB | Frontier Institute,<br>MSFR100410<br>(Miura et al., 2006) |
| DsRed | 1-225 aa<br>(GU253312) | Rb, Gp | AB_2571648<br>AB_2571647 | Tg | Frontier Institute,<br>MSFR101390<br>MSFR101410 |
| GABA<sub>A</sub>γ2 | 39-67 aa<br>(UniProt:<br>P22723) | Rb | AB_2571572 | KD | Synaptic systems<br>#224 003<br>(Rovo et al., 2014) |
| GFP | 1-238 aa<br>(YP_002302326) | Rb, Go | AB_2491093<br>AB_2571574 | Tg | Frontier Institute,<br>MSFR101890<br>MSFR101910<br>(Takasaki et al.,<br>2010) |
| GluA2 | 847-863 aa<br>(X57498) | Gp |  | KO | Yamasaki et al.,<br>2011 |
| GluN1 | 660-811 aa<br>(NM_000832.5) | Ms | AB_2571605 | IB | Millipore<br>(MAB363)<br>(Siegel et al.,<br>1994) |
| VGAT/VIAAT | 31-112 aa<br>(BC052020) | Rb | AB_2571622 | IB | Frontier Institute,<br>MSFR106120<br>(Miura et al., 2006) |
| VGLuT2 | 559-582 aa<br>(BC038375) | Gp | AB_2571621 | IB | Frontier Institute,<br>MSFR106280<br>(Miyazaki et al.,<br>2003) |

Antibody dilution for each experiment is described in the individual sections in Methods. Abbreviations: aa, amino acid residues; Go, goat polyclonal antibody; Gp,
guinea pig polyclonal antibody; IB, immunoblot with brain homogenates; KD, lack of immunolabeling in knockdown neurons; KO, lack of immunolabeling in knockout brains; Rb, rabbit polyclonal antibody; Tg, specific labeling in transgenic mouse brains.

